# Detection the Prevalence of Hepatitis C Virus among Iraqi People

**DOI:** 10.1101/2020.11.28.401968

**Authors:** Hasan Abd Ali Khudhair, Ali A. H. Albakaa, Khwam R. Hussein

## Abstract

Hepatitis C virus (HCV) infection is a major public health problem worldwide and remains a vital cause of chronic hepatitis. This study was aimed to detect the prevalence of HCV infection among Iraqi people. Four subjects of hemodialysis (HD) patients, thalassemia patients, blood donors and medical staff were enrolled in this study and evaluated for their serum anti-HCV-immunoglobulin G (IgG)-antibodies (Abs). The total frequency % of IgG anti-HCV Abs positivity was 3.2%, in which the highest frequency % was recorded among thalassemia patients followed by HD patients and then medical staff subjects, whereas the lowest frequency rate was reported within blood donors group. The frequencies of IgG anti-HCV Abs positivity were significantly elevated in males compared to females. For age groups, the results revealed higher infection rate of HCV among age group of 1-20 year followed by the age group of 21-40 year and then age group of 41-60 year, whereas the lowest rate of infection was recorded in age group >60 year. In conclusions, the prevalence rate of HCV infection among Iraqi people is similar to those in most of Asian and non-Asian studied populations and the infection rate was higher in males and inversely correlated with age of the patients. Blood transfusion, renal dialysis and health care workers (HCWs) were major sources of HCV infection. Thus, we recommend continuing surveillance of blood donors, HCWs and patients, in addition to HCV markers screening by molecular technique for the diagnosis of HCV during the window period in order to decrease the prevalence of HCV infection.

## INTRODUCTION

Hepatitis C virus is a small enveloped ribonucleic acid (RNA) virus belonging to the family Flaviviridae and genus hepacivirus. Hepatitis C virus genomic RNA was single-stranded with positive polarity, which was packaged by core protein and enveloped by a lipid bilayer containing two viral glycoproteins (E1 and E2) to form the virion. Despite the nucleotide sequence divergence among genotypes, all currently recognized HCV genotypes are hepatotropic and pathogenic (Li and Lo, 2015). Infection with HCV leads to an asymptomatic acute stage. However, approximately 75% of acutely infected patients face a substantial risk of developing chronic HCV infection. During the 2 decades after infection, 27% develop liver cirrhosis, and 25% develop hepatocellular carcinoma (HCC). Worldwide, an estimated 71 million people were living with chronic HCV infection (1% of the global population). Whilst, in the European Union/European Economic Area, it was estimated that more than 14 million people were living with chronic HCV infection, suggesting a relatively higher prevalence of 1.5% in this region (Han *et al*., 2019). Studies from regional countries showed that HCV prevalence was 1.1% in Yemen, less than 1% in Iran, 1.8% among young generation in Saudi Arabia, 4.0% among blood donors in Pakistan and 0.2% in Iraq (Hussein *et al*., 2017). In patients undergoing maintenance HD, the prevalence of HCV infection substantially increases and this disease has been shown to be associated with severe complications from chronic hepatitis to fatal cirrhosis and HCC (Khedmat *et al*., 2014). The natural history of HCV infection in dialysis patients remains incompletely understood; controversy continues even in patients with intact kidney function. Defining the natural history of HCV remains difficult for several reasons: the disease has a very long duration, it is mostly asymptomatic and determining its onset may be difficult (Fabrizi, 2013).

Hepatitis C virus is the most common cause of post-transfusion hepatitis (PTH) and end-stage liver disease in many countries. Regular blood transfusion in patients with hereditary hemolytic anemia, particularly thalassemia, has improved their overall survival, but carries a definite risk of acquisition of blood-borne virus infections, especially viral hepatitis (Boroujerdnia *et al*., 2009). Overall, the transmission of HCV involves direct exposure to contaminated blood and is associated with intravenous drug use, iatrogenic exposures, tattooing, body piercing, and less frequently through vertical transmission and high risk sexual behavior (Najim and Hassan, 2018). Blood transfusions contribute to the expanding transmission pool of viral infections, wherein even an asymptomatic person can transmit the infection. Screening and assessment of donors not only lessens the risk of transmission through infected blood products, but also gives an idea about the prevalence rates of the infections in the community. Evaluation and monitoring the prevalence and trend of HCV in blood donors is important for assessing quality and effectiveness of donor screening, public education, blood screening tests, and potential risk of transfusion-transmitted HCV infection (Khodabandehloo *et al*., 2013).

Regulation of screening tests together with the development and introduction of nucleic acid technique tests for HCV has improved blood safety. Despite advances in technology, transfusion-transmitted HCV infection still exists (Alhilfi *et al*., 2015). Health care workers wherever they work are at high risk of infectious blood-borne pathogens including HCV. The triggers related to their infection are being in contact with contaminated sharp instruments, injection malpractices, incorrect handling of biological materials, and insufficient education. The number of infected HCWs is often affected by the overall number of HCV infected population. This rate is often high in HCWs living in less developed countries (Abdelrheem *et al*., 2020). There is no vaccine exists to prevent HCV infection and treatment for HCV infection is costly. Thus, the prevention of primary HCV infection is very important. Any strategy to prevent HCV infection must be based on accurate data, including information about its prevalence. A few amounts of studies have been done on the prevalence of HCV infections in past years among Iraqi people in some Provinces. However, they are little and incomplete.

The diagnosis of HCV infection is based on the detection of anti-HCV Abs by enzyme linked immunosorbent assay (ELISA) and it is confirmation by a positive result obtained by an immunoblot assay or by the presence of HCV RNA. An improvised third-generation anti-HCV-ELISA with high sensitivity is widely used for patients screening. Most of the available third-generation ELISA tests for anti-HCV-Abs detection are based on either synthetic peptide antigens (Ags) alone or recombinant protein Ags or a combination of synthetic and recombinant protein Ags of HCV (Kesli, 2011).

## STUDY AIMS

The current study aimed to document the prevalence of HCV infection among HD patients, thalassemia patients, blood donors and medical staff Iraqi people. Such prevalence presumably might provide a help for prevention strategies of this infection and guide further researches.

## MATERIALS and METHODS

### Subjects

A total of 1650 individuals (1180 males and 470 females with an age range 1-85 year) attending the Public Health Laboratory, Hereditary Blood Diseases Center, Central Blood Bank and Renal Dialysis Unit/Al-Hussein Teaching Hospital at Thi-Qar province (Iraq) were enrolled in this study and they are classified into four groups; earliest one included 120 patients with renal failure, the second group included 220 patients suffering from thalassemia, the third group included 1259 blood donors subjects, whereas the last one included 51 subjects from the medical staff. A verbal consent was obtained from each subjects participating in this study to fulfill the international research ethical criteria. This study was carried out from August 2020 to October 2020.

### Data collection

Data of age, gender, residency and the result of laboratory test of anti-HCV-IgG-Abs were collected from records of the subjects at the mentioned above medical centers except the medical staff group which is underwent the following:

A. ***Samples collection and serum separation:*** From each subject; 3-4 ml of peripheral blood was collected by vein puncture. Blood samples were allowed to complete clotting processes at room temperature and then centrifuged at 1500 rpm for 10 minutes to get the sera that have been stored at ^-^20 C° until their needed for serological investigation.
B. ***Serological test:*** Serological test had been executed in Public Health Laboratory/Health Office of Thi-Qar. Anti-HCV-Abs (IgG) was detected by third generation ELISA kit from MyBioSource (United States of Americans (USA)). This assay employs the indirect qualitative enzyme immunoassay technique. The microtiter plate provided in this kit has been pre-coated with Ags (three recombinant proteins of the non-structural (NS) region (NS3 and NS4) and a peptide of the structural region). Samples are pipetted into the wells with anti-human immunoglobulin conjugated horseradish peroxidase. Any Abs present in the samples that are specific for the pre-coated Ag will bind to it. Following a wash to remove any unbound reagent, a substrate solution is added to the wells and color develops in proportion to the mount of human anti-HCV-Abs bound in the initial step. The color development is stopped and the intensity of color is measured at 450 nm.

### Statistical analysis

Statistical package for social sciences version 24 was used. Descriptive statistics included; the use of frequencies and relative frequencies. Chi-Square statistical test were used to test associations between the variables. The results being considered as statistically significant when the P-value was <0.05.

## RESULTS

Figure 1 showed the results of anti-HCV-(IgG)-Abs in total sum of the study groups. The findings revealed that out of 1650 subjects, only 53 (3.2%) were infected with HCV. In Figure 2, the results revealed the prevalence of HCV according to study groups. The highest prevalence was recorded among thalassemia patients 34/220 (15.4%) followed by HD patients 8/120 (6.6%) and then medical staff group 3/51 (5.8%), whereas the lowest prevalence was reported among blood donors group 8/1259 (0.63%).

**Figure (1):**
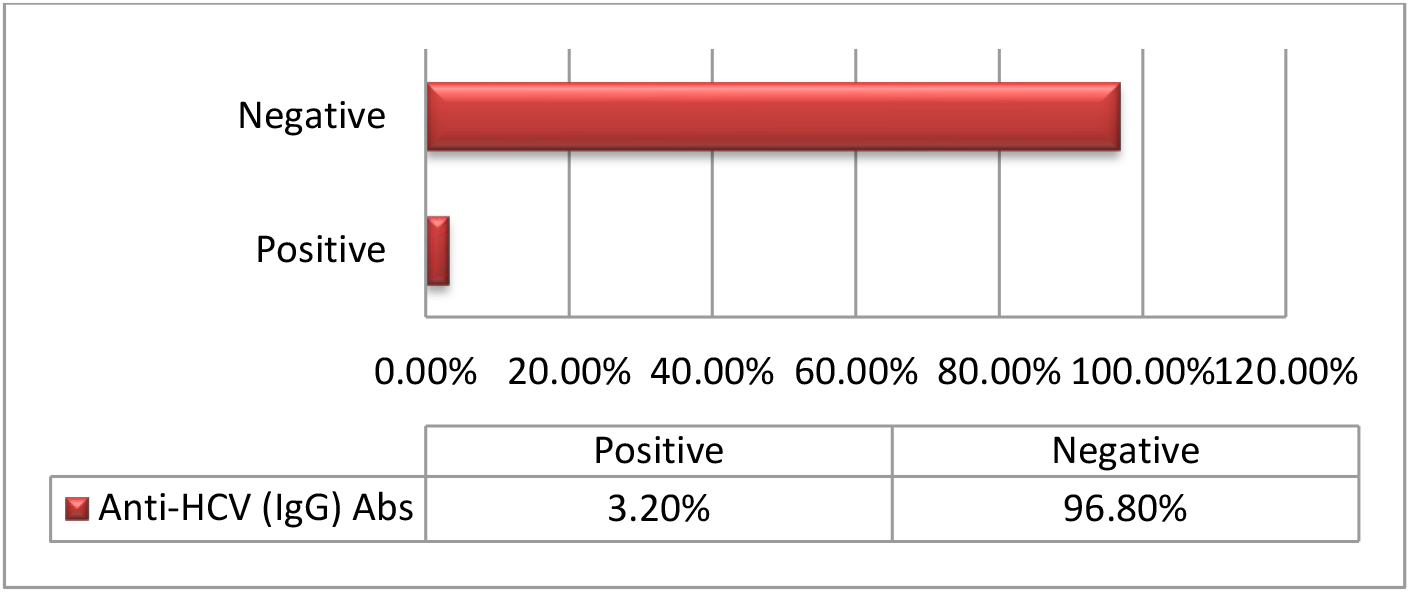
**Total prevalence of hepatitis C virus (HCV:** Hepatitis C virus, **IgG:** Immunoglobulin G and **Abs:** Antibodies).

**Figure (2):**
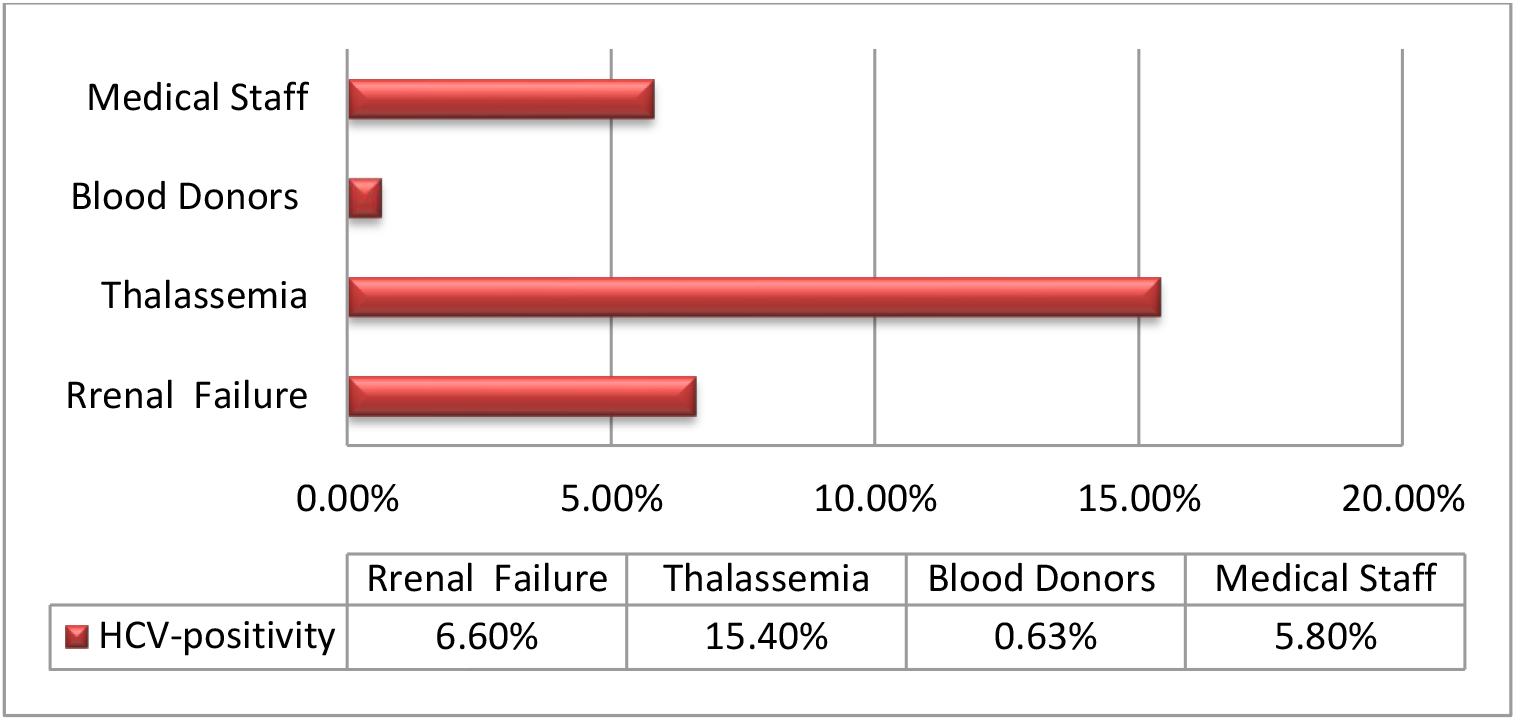
**The prevalence of hepatitis C virus according to the study groups (HCV:** Hepatitis C virus).

The findings of Table (1) were reported that infection rate of the HCV was higher among males 33/53 (62.3%) compared to females 20/53 (37.7%) with a significant differences *(p<0.05*) between them.

**Table (1):**
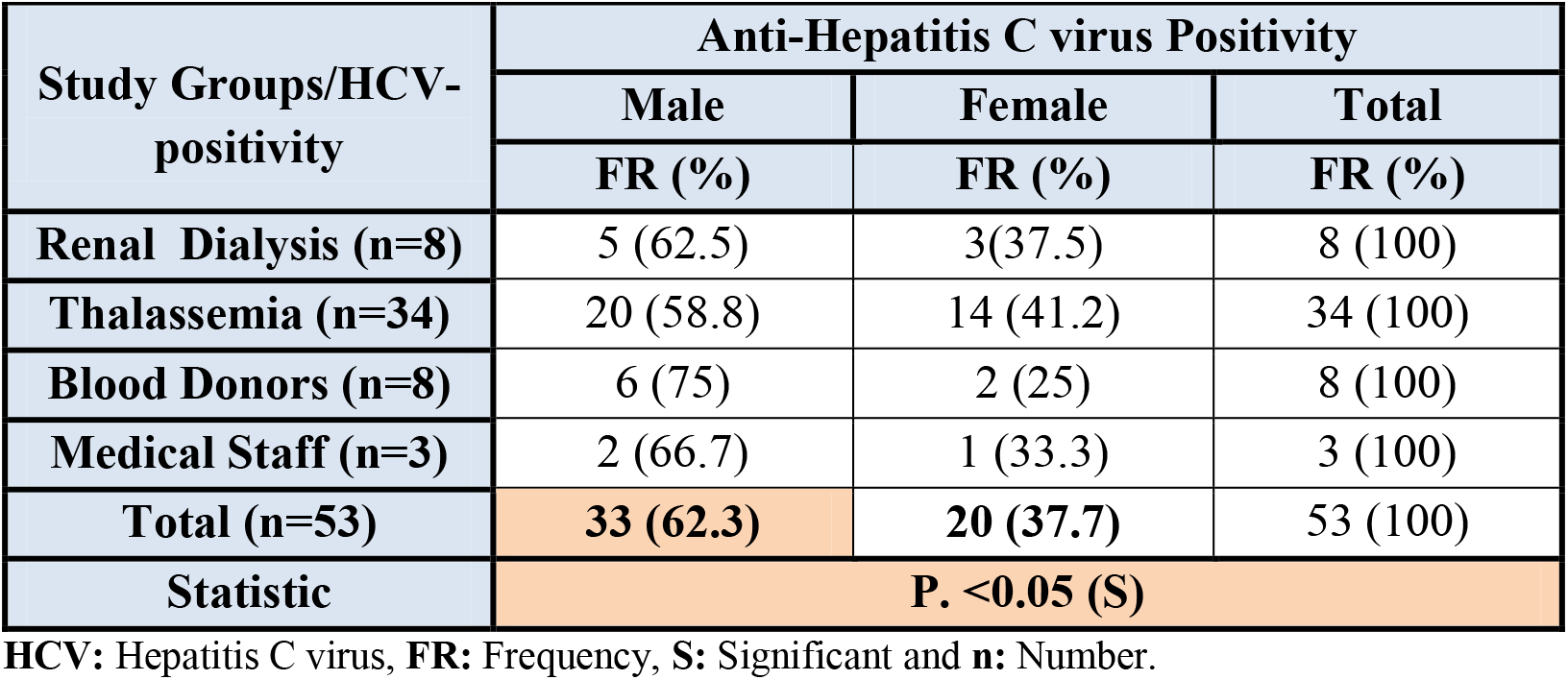
Prevalence of hepatitis C virus according to gender of the subjects.

| Study Groups/HCV-positivity | Anti-Hepatitis C virus Positivity |  |  |
| --- | --- | --- | --- |
|  | Male | Female | Total |
|  | FR (%) | FR (%) | FR (%) |
| Renal Dialysis (n=8) | 5 (62.5) | 3(37.5) | 8 (100) |
| Thalassemia (n=34) | 20 (58.8) | 14 (41.2) | 34 (100) |
| Blood Donors (n=8) | 6 (75) | 2 (25) | 8 (100) |
| Medical Staff (n=3) | 2 (66.7) | 1 (33.3) | 3 (100) |
| **Total (n=53)** | **33 (62.3)** | **20 (37.7)** | 53 (100) |
| **Statistic** | **P. <0.05 (S)** |  |  |

HCV: Hepatitis C virus, FR: Frequency, S: Significant and n: Number.

The results in Table (2) showed a significant different *(p<0.05)* in HCV infection rate between age groups of the study subjects. The higher infection percentages were 54.7% and 28.3% had been reported in age groups of 1-20 year and 21-40 year, respectively, followed by age group of 41-60 year that had infection percentage 13.2%, while the lower infection rate was recorded at the age group of more than 60 year, which was 3.8%.

**Table (2):** Prevalence of hepatitis C virus according to age of the subjects.

| Age/Study groups | Anti-Hepatitis C virus positive subjects |  |  |  |  |
| --- | --- | --- | --- | --- | --- |
|  | Renal Dialysis (n=8) | Thalassemia (n=34) | Blood Donors (n=8) | Medical Staff (n=3) | Total (n=53) |
|  | FR (%) | FR (%) | FR (%) | FR (%) | FR (%) |
| **1-20 Y** | 0 (0) | 29 (85.3) | 0 (0) | 0 (0) | 29 (54.7) |
| **21-40 Y** | 4 (50) | 5 (14.7) | 4 (50) | 2 (66.7) | 15 (28.3) |
| **41-60 Y** | 3 (37.5) | 0 (0) | 3 (37.5) | 1 (33.3) | 7 (13.2) |
| **>60 Y** | 1 (12.5) | 0 (0) | 1 (12.5) | 0 (0) | 2 (3.8) |
| **Total** | 8 (100) | 34 (100) | 8 (100) | 3 (100) | 53 (100) |
| **Statistic** | **P. <0.05 (S)** |  |  |  |  |

Y: year, FR: Frequency, S: Significant and n: Number.

## DISCUSSION

Hepatitis C virus infections are among the most prevalent infectious diseases in humans worldwide and it is associated with a broad range of clinical presentations ranging from acute or fulminant hepatitis to chronic infection that may be clinically asymptomatic or may progress to chronic hepatitis and liver cirrhosis. The prevalence of HCV infection varies from one country to another depending upon a complex mixture of behavioral, environmental and host factors (Tarky *et al*., 2013). The prevalence rate of HCV infection is decreasing in the developed countries; whereas, in developing countries such as Iraq, researchers still struggle to control the infection (Messina *et al*., 2015). However, the prevalence of HCV infection in the general population in Iraq is not available due to the unstable political and social situation for the past decades. To the best of our knowledge, the only published data on the prevalence of viral hepatitis in Iraq are from blood donor studies.

Different studies from neighbouring countries showed a variation in the prevalence rates of HCV ranging from 0.4% to 19.2% (Vriend *et al*., 2013). In this study the total prevalence of HCV infection was 3.2% (Figure 1). Consistent with our findings, another previous study at Duhok City (Iraq) found that the prevalence of HCV was 2.8% (Hussein *et al*., 2017). On other hand, the results of current study less than those in previous study conducted in Iraq by Hamied *et al*., (2010) that reported the HCV prevalence in Baghdad province (8.3%) and more than the findings obtained Tarky *et al*., (2013) in all over Iraqi governorates (0.4%) and Abdul-Kareem *et al*., (2001) in Al-Najaf province (0.34%). In comparison with results of the previous studies in other countries, the findings of the current study are in agreement with those in Turkey (2.4%)% (Erden *et al*., 2003), Thailand (2.8%) and Vietnam (2-2.9%), and less than those in Taiwan (4.4%), Pakistan (4.7%) and Egypt (14.9%), and more than those in Iran (0.5%), USA (0.01%), Australia (1.3%), China (1-1.9%), Saudi Arabia (1-1.9%) and Syria (1-1.9%) (Yen *et al*., 2003, Mostafa *et al*., 2010, Merat *et al*., 2010 and Sievert *et al*., 2011). These differences in the prevalence rate might be explained in part by the difference in study population, samples collection method and the techniques used in the diagnosis. Further population based studies are needed to explore this.

Patients with thalassemia and hemoglobinopathies need frequent blood transfusions which are important for improvement of their survival and reduce dangerous complications that produce from severe anemia. On other hand, this frequent blood transfusions will increase the probability of infection with different microbes especially; HCV, hepatitis B virus and human immunodeficiency virus (Rund and Rachmilewitz, 2005). Consistent with these findings, the prevalence ratio of HCV in our study was 15.4% (Figure 2) from 220 patients suffer from thalassemia. Previous studies in Duhok city (Al-Khaffaj and Al-Ghazal, 2019) and Mosul City-Iraq (Khalid and Abdullah, 2012) revealed that the prevalence of HCV among thalassemia patients were 11.05% and 17%, respectively, which in line with current data. The results of the current study was lower than other previous studies in other Iraqi cities including Diyala (26.4%) (Raham *et al*., 2011) and Karbala (37%) (Al-Greti, 2013), and higher than Babylon city (7.5%) (Tarish and Shakeer, 2014). These variation in the prevalence of HCV may be belong to variation in hygienic surveillance especially tests of blood. In addition, it reflects the variation of health awareness of the citizens in these cities. Hepatitis C virus characterized with low viral load, long incubation period which may be extended to six months and asymptomatic in acute and chronic periods, all these reasons will delay the seroconversion and finally, delay diagnosis of HCV in blood donors leading to increase the probability of HCV-infection among patients with thalassemia and hemoglobinopathies through hemolysis and this interprets the increasing of HCV-infection in thalassemia patients in our study. To reduce the HCV-infection, many countries insert the molecular biology technology within the routine protocol tests which recognize very low concentration of viral RNA (Whittaker *et al*., 2008).

In HD centers, HCV infection remains a major concern. Blood transfusion as well as nosocomial infection continues to play important roles in the transmission of HCV (Su *et al*., 2013). An overall prevalence of 6.6% of HCV infection was reported among HD patients in the Thi-Qar province according to the present study (Figure 2). This finding is closer to that reported in Baghdad province (7.1%) (Khattab, 2008) and Al-Anbar province (11.7%) (Al-Mashshhhadani, 2006). This prevalence is low when compared to reports from Sulaimani (Iraq) (26.7%) (Ramzi *et al*., 2010) and in other developing countries; (26%) in Hungary (Mendez-Sanchez *et al*., 2004), (24%) in Iran (Alavian *et al*., 2003), (30%) in India and (26%) in Oman (Alavian *et al*., 2003), but, it high when compared to reports from western countries; United Kingdom (0.4%) (Jadoul *et al*., 2004) and Mexico (1.2%) (Mendez-Sanchez *et al*., 2004).

As previously illustrated above HCV is a blood borne virus transmitted by the parenteral route. Infection frequently results in a chronic asymptomatic carrier state for many years before the development of symptomatic liver disease. Hepatitis C virus infected HCWs may therefore be unaware of their condition and their potential to infect patients. Healthcare workers, who perform exposure prone procedures, where injury to the workers may result in exposure of the patient’s open tissues to the blood of the workers, are theoretically at increased risk of infection with blood borne viruses (Thorburn *et al*., 2001). In the current study we found that the prevalence of HCV among HCWs was 5.8% (Figure 2). It is closer to the results of ALHaj *et al*., (2019) and Ansari and Dixit (2017) studies that conducted in Yemen (4.17%) and India (4%) among HCWs, respectively. However, it was more than the prevalence of HCV among HCWs in Dhaka-Bangladesh, Poland and India (1% and 1.9% and 3%, respectively) (Shah *et al*., 2017). This difference is because the HCWs adopt different preventive degree measures in different health-care centers. An overall, the present study may be helpful for understanding the prevalence of HCV among HCWs in Thi-Qar province (Iraq).

For the prevalence of HCV among blood donors, our study showed that the prevalence of HCV-Abs was 0.63%. In agreement with these findings, previous studies in Baghdad (Ataallah *et al*., 2011) and Babylon (Al-Juboury *et al*., 2010) governorates showed a closer prevalence rate of HCV infection (0.7% and 0.5%, respectively). On comparison with other countries, the prevalence of HCV among Kuwaiti national and non-Kuwaiti Arab first-time donors were found to be 0.8 and 5.4%, respectively (Ameen *et al*., 2005). In Jordan, a hospital-based study showed that the infection with HCV among blood donors was 0.9% (Al-Gani, 2011). Overall, the variations in the prevalence of HCV-infection may belong to different reasons such as sample size, type of technique used (ELISA, Minividas, Immunofluorescences or Chemiluminescence), variations in kit types and their trade mark, time of incubation during the test and the differences in blood test procedures between countries, cities and societies. In addition to that, the variations in customs and social customs in each society such as tattoo, body piercing and take drugs by injection. The level of hygienic surveillance as well as hygienic awareness of people may interpret all these variations in results. The results of the present study exhibited significantly higher frequency percent of HCV positivity among males compared to females (Table 2). The current study findings are consistent with previous findings that reported by others authors in Iraq (Abdul-Sada, 2011), Iran (Shakeri *et al*., 2013), Pakistan in blood donors, India in outpatients clinic visitors (Sood and Malvankar, 2010) and USA blood donors (Sheikh *et al*., 2013) as the prevalence was higher in males than females. In Egypt, higher HCV prevalence was detected in males compared to females among village residents (Sharaf-Eldin *et al*., 2007) and blood donors (Rushdy *et al*., 2009). In Pakistan, although the probability trends were slightly higher in males of all age groups than in females, the differences not statistically significant (Anwar *et al*., 2013). In a nationwide survey in Poland did not indicate a significant effect of gender on HCV prevalence (Hartleb *et al*., 2012). Considering the age groups distribution, the prevalence of HCV infection was significantly highest among persons with age group (1-20 year) and followed by age group (21-40 year) (Table 2), and these results were in agreement with Amin *et al*., (2004), that reported the 20-24 year old age group had the highest prevalence with strong majority of the infected population below the age of 50. In Western Pacific the pattern of seroprevalence across age in Australasia exhibits a rapid increase in prevalence peaking at 20-24 years age. In addition, in Central Europe an early peak in ages of 1-4 year is seen in Central Europe (Mohd *et al*., 2013). The prevalence of HCV infection showed a decrease with age after 60 years old, which may due to the decline in physical mobility and economic strength that leads to decrease the number of elderly examination.

## Conclusions

The prevalence rate of HCV infection among Iraqi people is 3.2%, which is similar to those in most of Asian and non-Asian studied populations and the infection rate was higher in males and inversely correlated with age of the patients. Blood transfusion, renal dialysis and HCWs were major sources of HCV infection. Thalassemic and HD patients are at risk of acquiring HCV infection. Therefore, blood donor screening protocol and effective screening techniques are likely to be needed to prevent speared of HCV infection among them. The risk of HCV infection is more among HCWs, hence HCWs should take proper precaution while handling blood. Aseptic procedures should be carried out to prevent needle stick injury.

In Iraq, information about the prevalence of HCV infections has generally been limited to laboratory data and personal interest of research projects in certain education institutes. Thus, we recommend continuing surveillance of blood donors, HCWs and patients, in addition to HCV markers screening by special type of molecular technique (polymerase chain reaction) for the diagnosis of HCV during the window period in order to decrease the prevalence of HCV-infection.

## Acknowledgements

We would like to thank the staff of all medical centers mentioned above for their infinite help and efforts during the data and samples collection. We also express our full heart gratitude to all the patients, volunteers and their families for their great assistance and cooperation. Special thanks to the staff of Training and Human Development Center/Health Office of Thi-Qar for their infinite help and efforts by facilitating the task of completing this research in coordination with the concerned medical centers.

## Notes

https://www.researchgate.net/publication/346427633_Detection_the_Prevalence_of_Hepatitis_C_Virus_among_Iraqi_People

### Competing Interest Statement

The authors have declared no competing interest.

